## Supplementary figures and images for "Surface exclusion of IncC conjugative plasmids and their relatives"

### Supplemental Figure 1

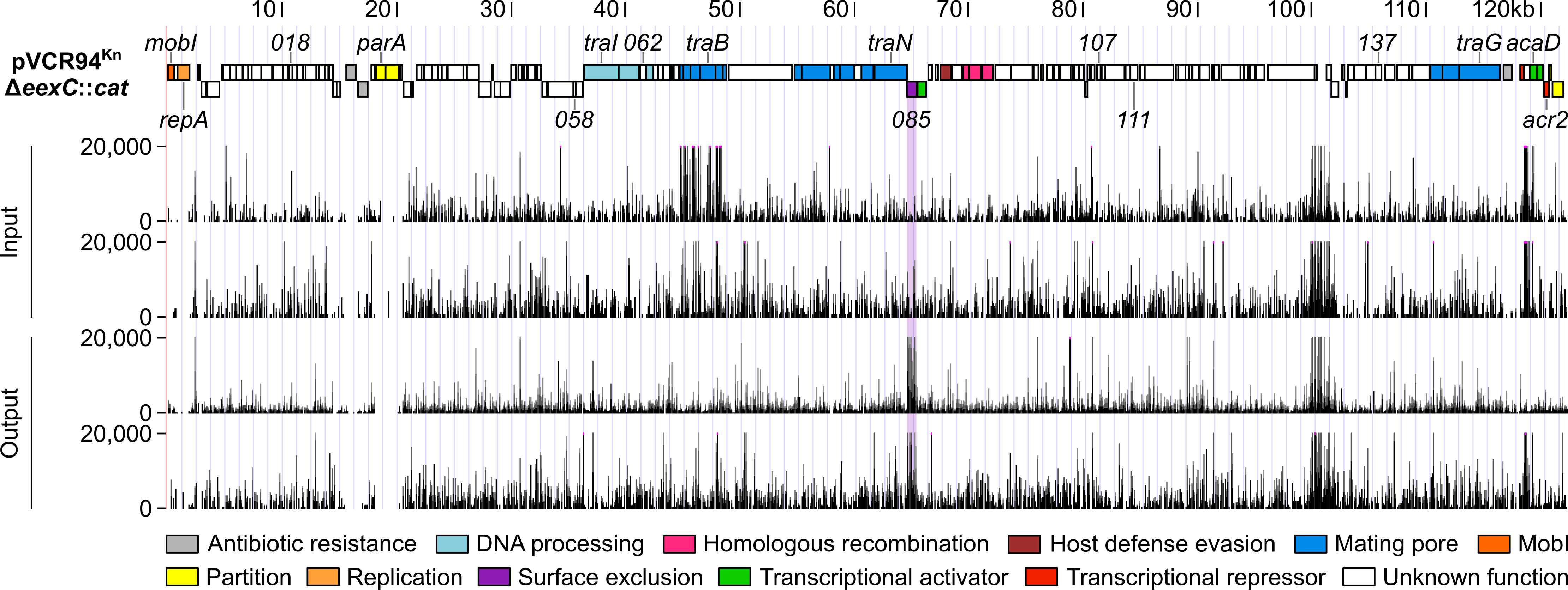

### Supplemental Figure 2

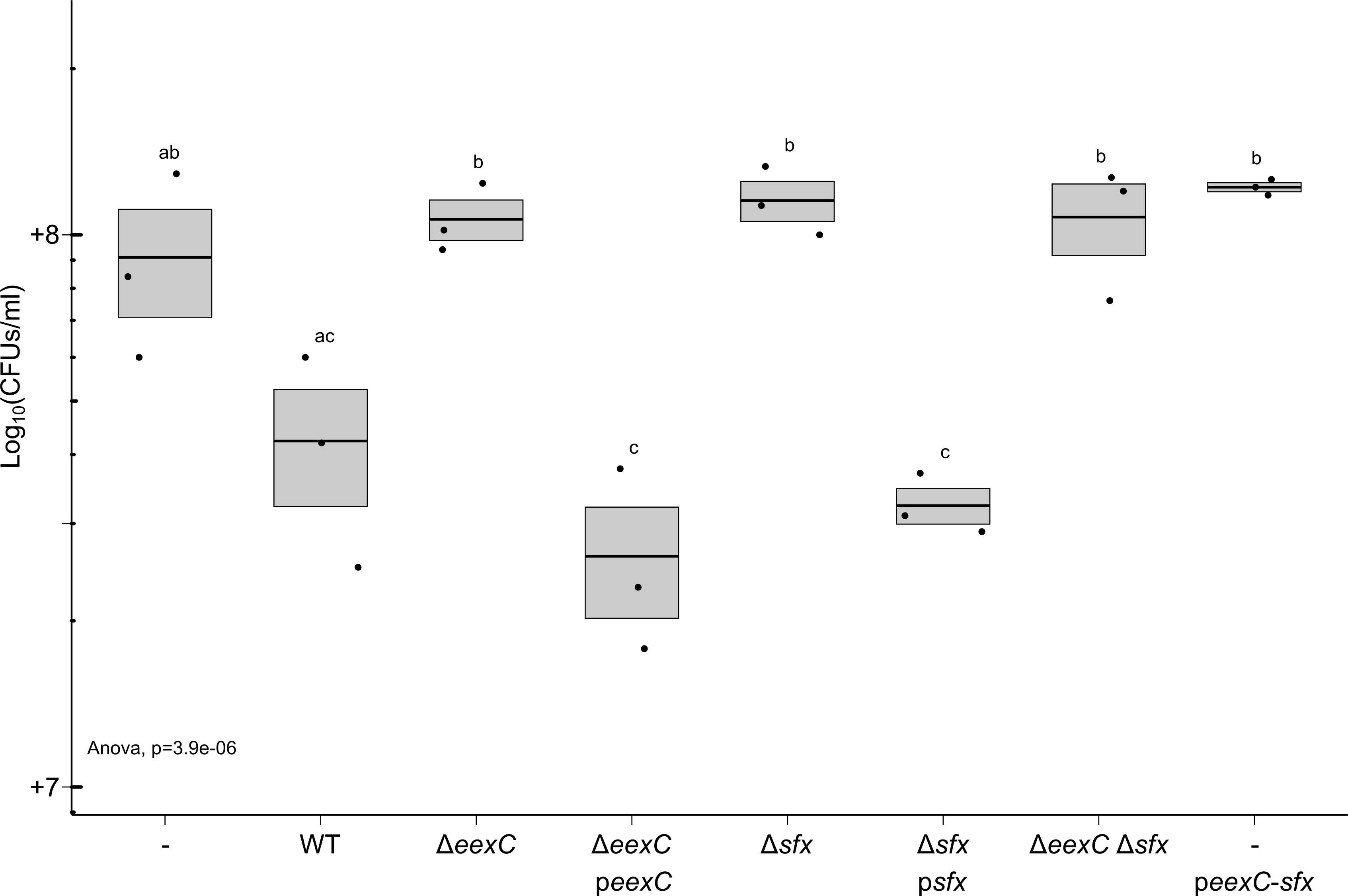

### Supplemental Figure 3

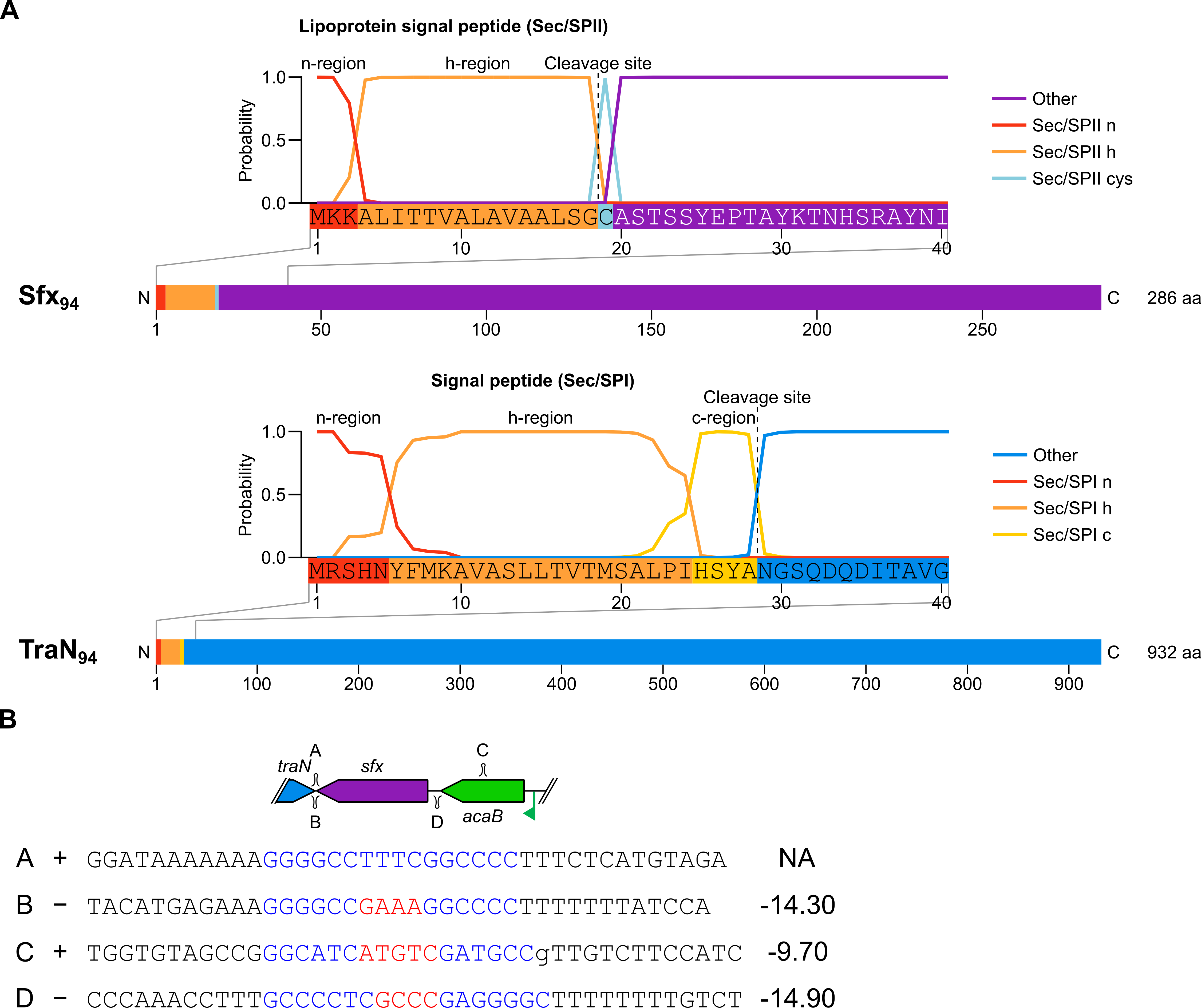

### Supplemental Figure 4

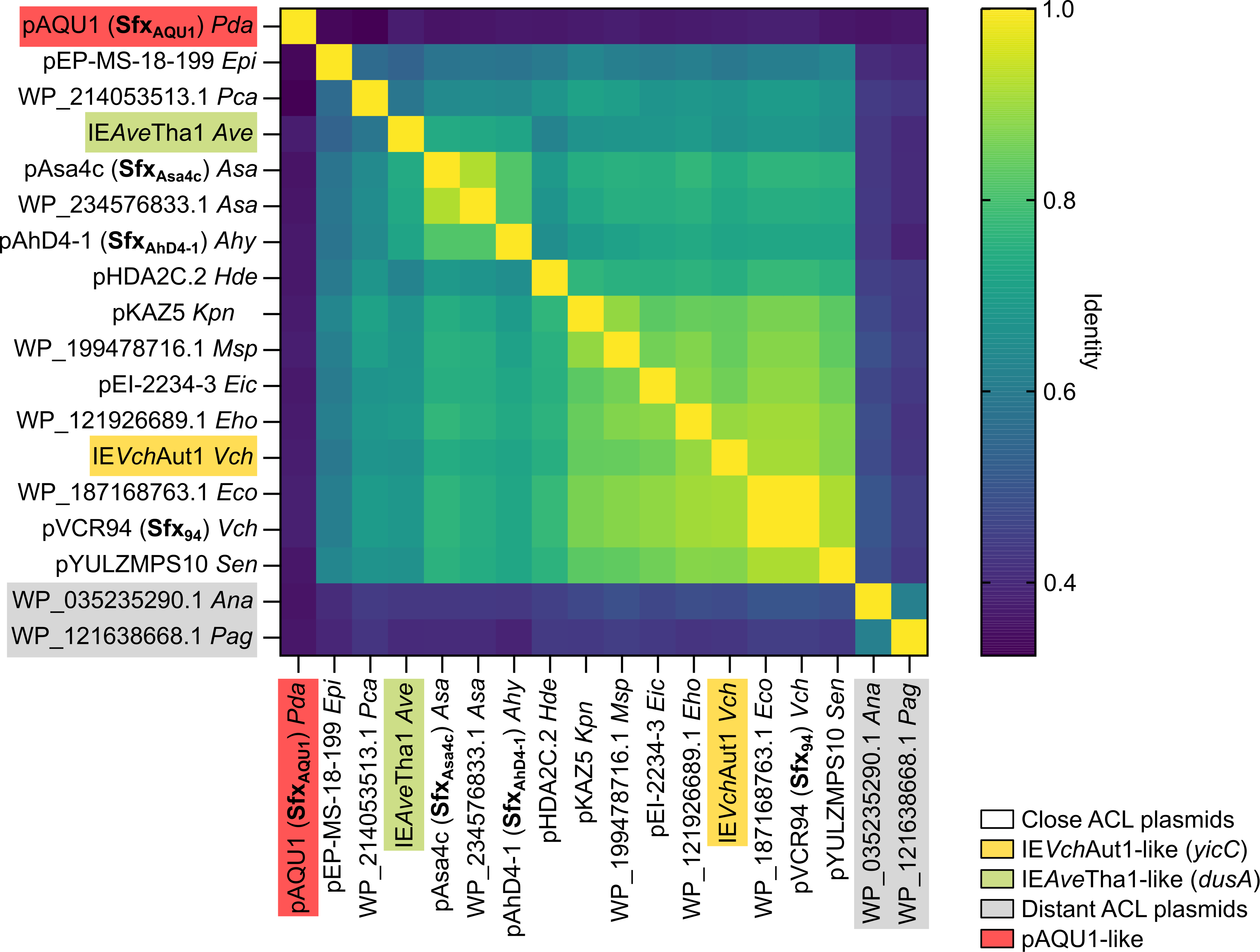

### Supplemental Figure 5

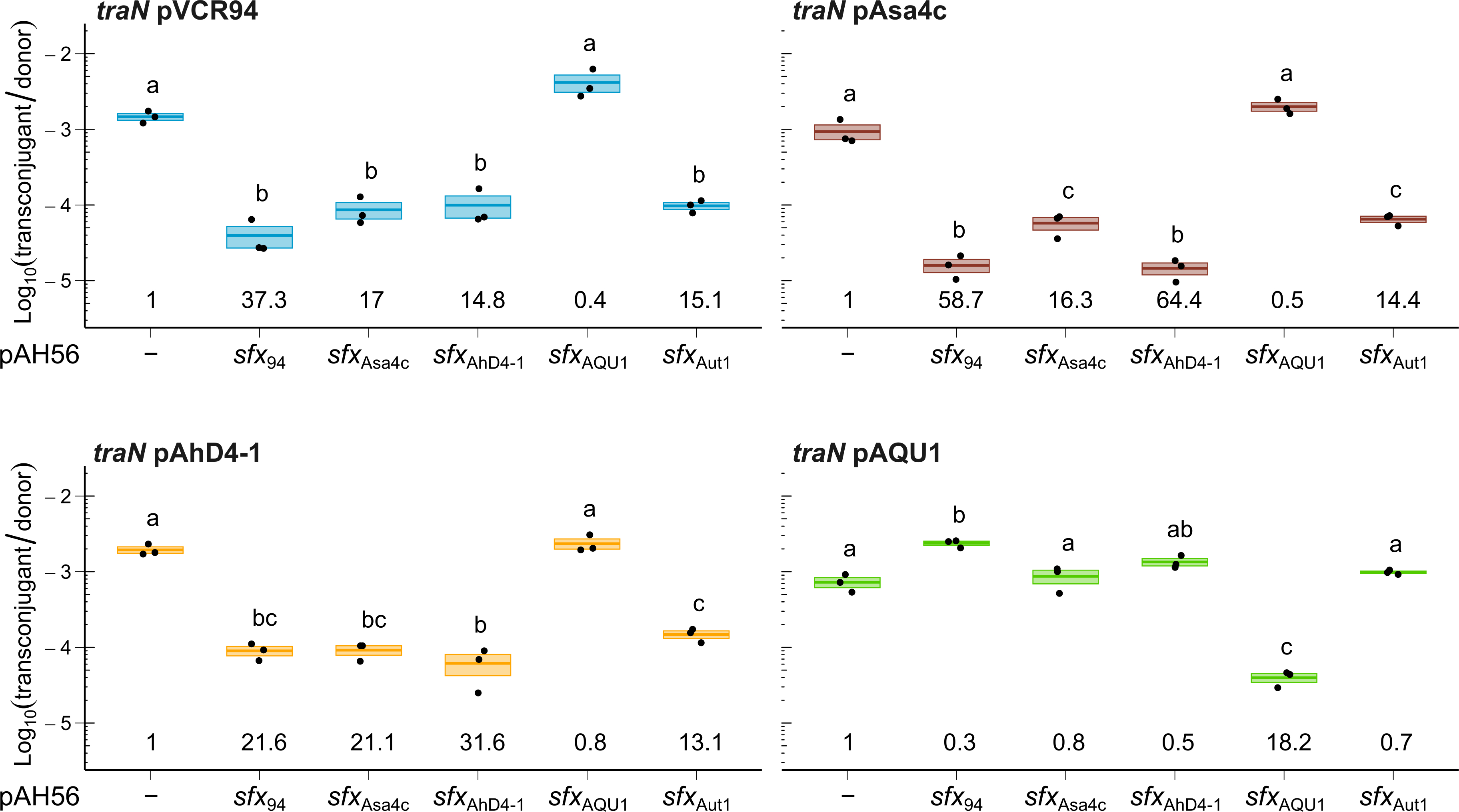
