## Supplemental Table 1 for "Surface exclusion of IncC conjugative plasmids and their relatives"

| **S1 Table. Primers used in this study** | |
| --- | --- |
| **Name** | **Nucleotide sequence (5' to 3')^a^** |
| 94delvcrx018.for | GCTCTCCGGGGCCTGATTTCACTATGAGGTGAATCTGTGTAGGCTGGAGCTGCTTCG |
| 94delvcrx019.rev | AACCTGGTTGGCGTTCGCGTTTTGAGTATTCGTCGTCATATGAATATCCTCCTTA |
| 94delvcrx062.for | AGGCCGTCTGCCCATATTGACCCAGTAAGAGCAGAGCTGTGTAGGCTGGAGCTGCTTCG |
| 94delvcrx062.rev | ACCAGCAGCCATGCGACCAGCAACCACGGCTTCTTCATCATATGAATATCCTCCTTA |
| 94delvcrx085.for | AAAAAGGGGCCTTTCGGCCCCTTTCTCATGTAGATTGAGTGTAGGCTGGAGCTGCTTCG |
| 94delvcrx085.rev | GTAAATCAGATTCAACTCAGGTAAGAACGGAGAACAACCATATGAATATCCTCCTTA |
| 94delvcrx107.for | GGCTGGTGGTTCCCTAACAAACATGGAGCCACAACTGTGTAGGCTGGAGCTGCTTCG |
| 94delvcrx108.rev | GAAGTAATGGCCCCGATGGGTTTAAAGTGGTTCTCACATATGAATATCCTCCTTA |
| 94delvcrx111.for | ACGGCTTGCATTGCTCTCTCGAAAGTGGAGAGTCAAGTGTAGGCTGGAGCTGCTTCG |
| 94delvcrx111.rev | GTACGCCGATTTCTCGGCGCACTCAGGTTGAAGCTACATATGAATATCCTCCTTA |
| 94delvcrx137.for | CTCCAGGAGCTTCAAAGCAAAGGTAATTTGGAATGAGTGTAGGCTGGAGCTGCTTCG |
| 94delvcrx137.rev | AGGACGACTTTTAGAGTGCCTAGATTTGCTCGATTACATATGAATATCCTCCTTA |
| 94deI145.for | ATGCCGTGGTAAACTGGAGGTAAAGAGTAAGGGGGTTGATGTGTAGGCTGGAGCTGCTTC |
| 94deI145.rev | ACCAACCAAATAATAAGGGGGCCAGCAGGCCCCCTTATTACATATGAATATCCTCCTTA |
| 94DelacaD.for | GCTATCTATCGCAACCTTCGTGATTTGTGAGGGGGGCGGAGTGTAGGCTGGAGCTGCTTC |
| 94DelacaC.rev | GCGCTCATCTTCTGGTCCGAAATGTCATAGTCTACTCATTACATATGAATATCCTCCTTA |
| 94delsfx.for | GTAAATCAGATTCAACTCAGGTAAGAACGGAGAACAACTACCTGTGACGGAAGATCAC |
| 94delsfx.rev | AAAAAGGGGCCTTTCGGCCCCTTTCTCATGTAGATTGATAGGAACTTCATTTAAATGG |
| 85RT | AGAAGGTAAGGGAAAGCAAC |
| sfxF | GGGTGAACGGACAAAGTAAG |
| sfxR | GTTTGCTAATAGCTGCGTAG |
| 086F2 | AGAAGCACAAGCAGCAGTG |
| 086R2 | TGGAAATGTGTTCCAGGATG |
| 94sfxNdeI.f | GATCCATATGAAAAAGGCTTTAATCACCA |
| 94sfxSalI.r | GATCGTCGACTTACGCTTCTGGGTAAAC |
| Asa4sfxNdeI.f | GATCCATATGAATAAGATACTGATCGCCAC |
| Asa4sfxSalI.r | GATCGTCGACTCAAGAAGGGTAAACAAAGAG |
| AhD4sfxNdeI.f | GATCCATATGAACAAGTCTCTGATCGCCAC |
| AhD4sfxSalI.r | GATCGTCGACTCATGAAGGATAAACAAAGAG |
| AQU1sfxNdeI.f | GATCCATATGATAAATGGAAAACTAATAGG |
| AQU1sfxSalI.r | GATCGTCGACTTAGTTCTCTGGGTGTATG |
| Aut1sfxNdeI.f | gatcCATATGAAAAAGGCTTTAATCACCACC |
| Aut1sfxSalI.r | gatcGTCGACTCAACCTTCCGGATAGGCAAAG |
| eexCNdeI.f | GATCCATATGAAACATGTGGTCAATATTCTTC |
| eexCSalI.r | GATCGTCGACTTATTCGTCTCCAGCTCCAA |
| pAH56_3xFlag.F | CATGACATCGACTACAAGGATGACGATGACAAGTAAGTCGACGGATCCCCGGAATT |
| 94sfx_3xFlag.R | ATCTTTATAATCACCGTCATGGTCTTTGTAGTCCGCTTCTGGGTAAACAAATA |
| 94del86acaB.for | TGAGGCGTTGCTGCCATTGGTAAGTGGCGTCGGTGCGTGTAGGCTGGAGCTGCTTC |
| 94del144traG.rev | TAAGGGGGCCTGCTGGCCCCCTTATTATTTGGTTGGTCTTTTATTACATATGAATATCCTCCTTA |
| 94eexBamHI.for | NNNGGATCCTTAAACTGCGTTGTTAGCCA |
| 94sfxHindIII.rev | NNNAAGCTTCAGCATAGACCCTCAAACAG |
| Asa4traNEcoRI.f | GTCAGAATTCAAGGAGGATAATAA ATGAACAACCAATCGGTA |
| Asa4traNSalI.r | GTCAGTCGACTTAATTGCTACCAGTAGGA |
| AhD4traNEcoRI.for | NNNNGAATTCAAGGAGGAATAATAA ATGAACAATCAATCGGTAG |
| AhD4traNEcoRI.rev | NNNNGAATTCTTATGGCTGCCTAGCTCCTG |
| AQU1traNEcoRI.for | NNNNGAATTCAAGGAGGAATAATAA ATGAGAAATTATACAAGGTTG |
| AQU1traNEcoRI.rev | NNNNGAATTCTTAATAACCTGGAGCACCTG |
| 94sfx-lacZ.f | TCAACTCAGGTAAGAACGGAGAACAACATGAAAAAGCTGGCCGTCGTTTTACAACGTCG |
| 94sfx-lacZ.r | GGGGCCTTTCGGCCCCTTTCTCATGTAGATTGATTAGCAGCATTACACGTCTTGAG |
| 94eex-lacZ.f | CTGGAGGTAAAGAGTAAGGGGGTTGATATGAAACATCTGGCCGTCGTTTTACAACGTCG |
| 94eex-lacZ.r | CCAACCAAATAATAAGGGGGCCAGCAGGCCCCCTTAGCAGCATTACACGTCTTGAG |
| 94traN-lacZ.f | CATGGGAAGGTTGAAATGGAGAACACAATGCGAAGTCTGGCCGTCGTTTTACAACGTCG |
| 94traN-lacZ.r | CTACATGAGAAAGGGGCCGAAAGGCCCCTTTTTTTAGCAGCATTACACGTCTTGAG |
| ^a^restriction sites are underlined | |
