## Supplemental Table 2 for "Surface exclusion of IncC conjugative plasmids and their relatives"

| **S2 Table. Predicted AcaCD binding sites** | | | | | | | | | | |
| --- | --- | --- | --- | --- | --- | --- | --- | --- | --- | --- |
| **Sequence** | **Strand** | | **Start^1^** | | **End^1^** | | **p-value** | | **q-value** | **Matched Sequence** |
| IE*Vch*Aut1 | - | | 41111 | | 41138 | | 2.71E-08 | | 0.00119 | GATTTGTCCAAAATGGGCACTTTGGCGT |
| IE*Vch*Aut1 | + | | 48422 | | 48449 | | 4.25E-08 | | 0.00119 | GATTTGTCCAAATTGGGCAGTTTGGCGT |
| IE*Vch*Aut1 | - | | 44366 | | 44393 | | 5.49E-08 | | 0.00119 | GTTTTGTCCAAATTGGGCAGTTTGGCGT |
| IE*Vch*Aut1 | + | | 59122 | | 59149 | | 7.05E-08 | | 0.00119 | GTTTTGTCCAAATTGGGCACTTTGGCGT |
| IE*Vch*Aut1 | + | | 40239 | | 40266 | | 1.76E-07 | | 0.00223 | AAACTGCCCAATACGGACAGATTGAGCG |
| IE*Vch*Aut1 | - | | 40234 | | 40261 | | 1.98E-07 | | 0.00223 | AATCTGTCCGTATTGGGCAGTTTGAGAG |
| IE*Vch*Aut1 | + | | 54633 | | 54660 | | 6.56E-07 | | 0.00569 | CTATTGTCCAAAAAGGGCAGTTTGGCGT |
| IE*Vch*Aut1 | - | | 48417 | | 48444 | | 6.73E-07 | | 0.00569 | AAACTGCCCAATTTGGACAAATCGATCA |
| IE*Vch*Aut1 | + | | 41116 | | 41143 | | 2.01E-06 | | 0.0151 | AAAGTGCCCATTTTGGACAAATCGGTCA |
| IE*Vch*Aut1 | + | | 44371 | | 44398 | | 5.22E-06 | | 0.0353 | AAACTGCCCAATTTGGACAAAACAGTAA |
| IE*Vch*Aut1 | - | | 62638 | | 62665 | | 1.10E-05 | | 0.0673 | CATTTGTCCAAATCGGGCAGTTTGGCGT |
| IE*Vch*Aut1 | + | | 51190 | | 51217 | | 1.48E-05 | | 0.0835 | GTTTTGCCAAAATTTGGGGCGTGGAGAG |
| IE*Vch*Aut1 | + | | 53112 | | 53139 | | 4.89E-05 | | 0.254 | TGATCGCCCAAGTAGAGAGAATAGACAG |
| IE*Vch*Aut1 | - | | 49502 | | 49529 | | 5.59E-05 | | 0.27 | GTCACGCCCATCTAGGGCAACACAGGTT |
| IE*Vch*Aut1 | - | | 60712 | | 60739 | | 8.09E-05 | | 0.365 | TGGATGACCTCAAAGGAGACTTAGCGGA |
| IE*Vch*Aut1 | + | | 55315 | | 55342 | | 9.37E-05 | | 0.388 | GGTGAACCACAATACAGCCGCTCCACAA |
| IE*Vch*Aut1 | - | | 41120 | | 41147 | | 9.76E-05 | | 0.388 | AAAGTGACCGATTTGTCCAAAATGGGCA |
| IE*Sch*Chn1 | - | | 5111 | | 5138 | | 8.51E-12 | | 5.18E-07 | TTTGCGCCCTAAAAGGGCAAGTACAGAG |
| IE*Sch*Chn1 | - | | 8386 | | 8413 | | 3.00E-08 | | 0.000647 | ATTTTGTCCAAAAAGGGCAGTTTGGCGT |
| IE*Sch*Chn1 | + | | 20814 | | 20841 | | 4.01E-08 | | 0.000647 | TTTTTGTCCAAATTGGGCACTTTGGCGT |
| IE*Sch*Chn1 | + | | 12396 | | 12423 | | 4.25E-08 | | 0.000647 | GATTTGTCCAAATTGGGCAGTTTGGCGT |
| IE*Sch*Chn1 | - | | 2645 | | 2672 | | 9.54E-08 | | 0.00116 | AATGTGTCCGTATTGGGCAGTTTGAGAG |
| IE*Sch*Chn1 | + | | 2650 | | 2677 | | 2.22E-07 | | 0.00225 | AAACTGCCCAATACGGACACATTGAGCG |
| IE*Sch*Chn1 | + | | 19163 | | 19190 | | 6.56E-07 | | 0.00513 | CTATTGTCCAAAAAGGGCAGTTTGGCGT |
| IE*Sch*Chn1 | - | | 12391 | | 12418 | | 6.73E-07 | | 0.00513 | AAACTGCCCAATTTGGACAAATCGATCA |
| IE*Sch*Chn1 | - | | 20809 | | 20836 | | 1.66E-06 | | 0.0112 | AAAGTGCCCAATTTGGACAAAAAAGAAA |
| IE*Sch*Chn1 | + | | 5116 | | 5143 | | 3.52E-06 | | 0.0215 | TACTTGCCCTTTTAGGGCGCAAAAAGTG |
| IE*Sch*Chn1 | - | | 24583 | | 24610 | | 1.10E-05 | | 0.0607 | CATTTGTCCAAATCGGGCAGTTTGGCGT |
| IE*Sch*Chn1 | + | | 15720 | | 15747 | | 1.48E-05 | | 0.0752 | GTTTTGCCAAAATTTGGGGCGTGGAGAG |
| IE*Sch*Chn1 | + | | 8391 | | 8418 | | 2.51E-05 | | 0.118 | AAACTGCCCTTTTTGGACAAAATGGAAA |
| IE*Sch*Chn1 | - | | 3818 | | 3845 | | 2.88E-05 | | 0.124 | TATCTTCCCAAGTTGGGGGGGTTCACAT |
| IE*Sch*Chn1 | + | | 22345 | | 22372 | | 3.05E-05 | | 0.124 | TTAGAACCCTACAAGGGCGAAAAACTCT |
| IE*Sch*Chn1 | + | | 20887 | | 20914 | | 3.55E-05 | | 0.134 | GTAGAGATAAATGAGGACACTATGGCGG |
| IE*Sch*Chn1 | - | | 14029 | | 14056 | | 3.75E-05 | | 0.134 | GTCGCGCCCATCTAGGGCAACACAGGTT |
| IE*Sch*Chn1 | - | | 6383 | | 6410 | | 4.93E-05 | | 0.167 | AAAGCAAACTAATTGGACACTTAGCCTA |
| IE*Sch*Chn1 | - | | 22657 | | 22684 | | 6.74E-05 | | 0.216 | TGGATGACCTAAAAGGAGACTTAGCAGA |
| IE*Sch*Chn1 | - | | 19158 | | 19185 | | 7.23E-05 | | 0.22 | AAACTGCCCTTTTTGGACAATAGACGGC |
| IE*Spu*Chn1 | - | | 1276443 | | 1276470 | | 2.71E-08 | | 0.000762 | GATTTGTCCAAAATGGGCACTTTGGCGT |
| IE*Spu*Chn1 | - | | 1279691 | | 1279718 | | 3.00E-08 | | 0.000762 | ATTTTGTCCAAAAAGGGCAGTTTGGCGT |
| IE*Spu*Chn1 | + | | 1283885 | | 1283912 | | 4.25E-08 | | 0.000762 | GATTTGTCCAAATTGGGCAGTTTGGCGT |
| IE*Spu*Chn1 | + | | 1275581 | | 1275608 | | 8.18E-08 | | 0.00101 | AAACTGCCCAATACGGACAATTTGAGCG |
| IE*Spu*Chn1 | + | | 1291938 | | 1291965 | | 9.42E-08 | | 0.00101 | TTTTTGTCCAAATTGGGCACTTTGCCGT |
| IE*Spu*Chn1 | - | | 1275576 | | 1275603 | | 1.76E-07 | | 0.00157 | AAATTGTCCGTATTGGGCAGTTTGAGAG |
| IE*Spu*Chn1 | + | | 1290269 | | 1290296 | | 9.81E-07 | | 0.00753 | CTATTGTCCCAAAAGGGCAGTTTGGCGT |
| IE*Spu*Chn1 | - | | 1291933 | | 1291960 | | 1.66E-06 | | 0.0112 | AAAGTGCCCAATTTGGACAAAAAAGAAA |
| IE*Spu*Chn1 | + | | 1276448 | | 1276475 | | 2.01E-06 | | 0.012 | AAAGTGCCCATTTTGGACAAATCGGTCA |
| IE*Spu*Chn1 | - | | 1283880 | | 1283907 | | 2.65E-06 | | 0.0142 | AAACTGCCCAATTTGGACAAATCAGAAA |
| IE*Spu*Chn1 | - | | 1295723 | | 1295750 | | 1.10E-05 | | 0.0536 | CATTTGTCCAAATCGGGCAGTTTGGCGT |
| IE*Spu*Chn1 | + | | 1298157 | | 1298184 | | 1.44E-05 | | 0.0613 | TTGCCGCCGATAATGGTCACTTCGCCGT |
| IE*Spu*Chn1 | + | | 1286847 | | 1286874 | | 1.48E-05 | | 0.0613 | GTTTTGCCAAAATTTGGGGCGTGGAGAG |
| IE*Spu*Chn1 | + | | 1279696 | | 1279723 | | 2.51E-05 | | 0.0964 | AAACTGCCCTTTTTGGACAAAATGGAAA |
| IE*Spu*Chn1 | + | | 1292011 | | 1292038 | | 3.55E-05 | | 0.127 | GTAGAGATAAATGAGGACACTATGGCGG |
| IE*Spu*Chn1 | - | | 1277688 | | 1277715 | | 4.93E-05 | | 0.166 | AAAGCAAACTAATTGGACACTTAGCCTA |
| IE*Spu*Chn1 | + | | 1296706 | | 1296733 | | 8.83E-05 | | 0.276 | CTAGTACCAAGAATAGACAGCACAACAA |
| IE*Spu*Chn1 | + | | 1290951 | | 1290978 | | 9.37E-05 | | 0.276 | GGTGAACCACAATACAGCCGCTCCACAA |
| IE*Spu*Chn1 | - | | 1276452 | | 1276479 | | 9.76E-05 | | 0.276 | AAAGTGACCGATTTGTCCAAAATGGGCA |
| IE*Ave*Tha1 | + | | 50541 | | 50568 | | 3.08E-11 | | 1.07E-06 | AAGGTGCCCAAAAAGGGCAGATCCAGAG |
| IE*Ave*Tha1 | - | | 50536 | | 50563 | | 3.24E-08 | | 0.000564 | GATCTGCCCTTTTTGGGCACCTTGAGCG |
| IE*Ave*Tha1 | + | | 53641 | | 53668 | | 1.64E-07 | | 0.00191 | GCTTTGCCCGAAAAAGGCACTTCCAGCG |
| IE*Ave*Tha1 | + | | 56024 | | 56051 | | 7.29E-05 | | 0.617 | AAACAGCTCAAGGAAGGCACATAGGCAG |
| IE*Ave*Tha1 | + | | 50814 | | 50841 | | 8.88E-05 | | 0.617 | TCTGCGACCATTTTTGGCAACACAATCG |
| ^1^Start and end positions refer to the chromosome or contig sequences available on Genbank:  IE*Vch*Aut1, NZ_VIPE01000140.1; IE*Sch*Chn1, NZ_JAIUZX010000023.1; IE*Spu*Chn1,  NZ_CP104755.1; IE*Ave*Tha1, JAAQQQ010000037.1. | | | | | | | | | | |
